# Multiple introductions shaped genomic diversity and demographic history of invasive box tree moth (*Cydalima perspectalis*) populations in North America

**DOI:** 10.64898/2026.09.16.751060

**Authors:** Aarati Basnet, Tanisha Chaudhary, Jennifer Antonides, Cody LaDuke, M. Lukas Seehausen, He Yin, Sarah Eichler, Yunke Wu, Sangeet Lamichhaney

## Abstract

The box tree moth *(Cydalima perspectalis)* is an invasive pest of boxwood plants *(Buxus spp.)* that has rapidly expanded across Europe and, since its first detection in Canada in 2018, across North America. Understanding its introduction pathways and early population structure is critical for effective surveillance and management. We integrated genome-wide single nucleotide polymorphism (SNP), mitochondrial cytochrome c oxidase I *(COI)*, and morphological data to investigate the early invasion history of *C. perspectalis* across North America. Genome-wide analyses revealed pronounced genetic differentiation between North American and European populations, together with substantial genetic structure within North America. Delaware was genetically distinct from other North American populations and showed greater affinity with European populations, consistent with a separate introduction history. In contrast, New York, Massachusetts, Michigan, Ohio, and Ontario showed greater genetic similarity, suggesting shared ancestry and/or regional connectivity, whereas Virginia and West Virginia exhibited additional differentiation. Mitochondrial data supported shared ancestry among most North American populations, while Delaware shared a haplotype group with the Croatian European population. D-statistic analyses identified asymmetric allele sharing among several North American and European populations, consistent with multiple introductions and subsequent admixture. Morphometric analyses revealed significant geographic variation in several body and wing traits. Together, these complementary datasets indicate that multiple introductions, regional genetic structuring, and admixture have shaped the North American invasion. These findings highlight the value of integrating genomic approaches with nursery and border surveillance to identify introduction pathways and inform targeted containment and long-term management of this emerging horticultural pest.

**Key Message:** • The invasion history of box tree moth in North America remains poorly understood.

• Genomic, mitochondrial, and morphological data revealed multiple invasion pathways.

• Delaware showed a distinct genetic history from most North American populations.

• Genetic structure revealed regional differences among North American populations.

• These findings can improve surveillance and targeted containment of this pest.

## Introduction

Invasive alien species (IAS) are major threats to biodiversity and ecosystem functioning (Early et al. 2016). Although natural invasions have occurred episodically throughout evolutionary history and have generally been limited in geographic scale and scope (Vermeij 1991), human-mediated invasions are global, ongoing, and occurring at rates far exceeding those expected from natural dispersal (Ricciardi 2007). Contemporary biological invasions affect virtually all continents and oceans, including remote and environmentally challenging regions, and are increasingly facilitated by human activities. International trade and transportation are among the primary drivers of biological invasions, while climate change, socio-economic development, migration, and tourism can further influence the establishment and spread of non-native species (Essl et al. 2020). IAS can produce direct and indirect cascading effects on ecological communities (Tobin 2018), posing substantial threats to agriculture, biodiversity, human livelihoods, and economies (Andersen et al. 2004; McGaughran et al. 2024).

In the United States, the economic costs associated with IAS have been estimated at approximately $120 billion USD annually, including nearly $40 billion USD in annual losses to crop and forest productivity attributed to invasive insects and pathogens (Pimentel et al. 2005; Paini et al. 2016). The movement of live plants is an important pathway for the introduction of phytophagous insects (Yamanaka et al. 2015), as arthropods can be transported unintentionally as contaminants of horticultural, agricultural, and forestry products through international trade and other human-mediated transportation networks (Hulme et al. 2008; Hulme 2009). Plant trade is estimated to have been responsible for the introduction of approximately 70% of invasive insects and pathogens detected in the United States between 1860 and 2006 (Liebhold et al. 2012). The introduction and subsequent spread of the Japanese beetle *(Popillia japonica)*, for example, has been linked to the importation of ornamental plants into the United States in the early twentieth century (Fleming 1963; Althoff and Rice 2022). Such examples illustrate how movement of plants can create opportunities for repeated introductions and long-distance establishment of herbivorous pests.

Prevention is generally considered the most cost-effective approach to managing IAS, followed by eradication through early detection and rapid response, containment, and prevention of further spread (Geburzi and McCarthy 2018). Once an invasive species becomes widely established, complete eradication can become increasingly difficult, requiring sustained efforts to contain populations and reduce dispersal into new areas (Harvey and Mazzotti 2014). The recent establishment of the invasive box tree moth *(Cydalima perspectalis)* in North America provides a suitable example of an emerging horticultural pest for which understanding introduction pathways during the early phase of invasion may inform surveillance and management. *C. perspectalis* is native to East Asia, including China (Xiao et al. 2011), Korea (Kim and Park 2013), Japan (Maruyama and Shinkaji 1987), India (Hampson 1896), Pakistan (Sial et al. 2017), and Far East Russia (Musolin et al. 2022) **(Fig. 1)**. The species was first detected in Europe in Germany in 2007 (Bras et al. 2019a) and subsequently spread rapidly across the continent. It has now been reported from more than 30 Eurasian countries (Bras et al. 2019b), where it threatens both cultivated and natural boxwood *(Buxus)* plant species (Mitchell et al. 2018). Its rapid expansion in Europe has been attributed in part to the widespread availability of *Buxus* plant hosts and delays in detection following its initial establishment (Coyle et al. 2022). In North America, by contrast, the invasion is comparatively recent and remains at an earlier stage of establishment in many regions. The species was first detected in Ontario, Canada, in 2018 (Wiesner et al. 2021) and subsequently detected in New York, USA, in 2021 (Proxmire 2022). Since then, *C. perspectalis* has been reported from multiple U.S. states, including Michigan (2022), Massachusetts (2023), Ohio (2023), Delaware (2024) (Boggs et al. 2025), Pennsylvania (2024) (Prade 2025), Maryland (2025) (MDA 2025), Virginia (2025) (VDACS 2025), and West Virginia (2025) (WVDA 2025). Larvae feed on leaves and shoots of *Buxus* species and can cause severe defoliation, repeated damage, and ultimately plant mortality (Wan et al. 2014) **(Fig. 1A-C)**. The economic consequences are potentially substantial because approximately 12 million boxwood plants valued at $140 million are sold annually in the United States (NASS 2020), making *C. perspectalis* an important emerging threat to the North American nursery and ornamental horticulture industry (Dhakal et al. 2022).

**Figure 1.**
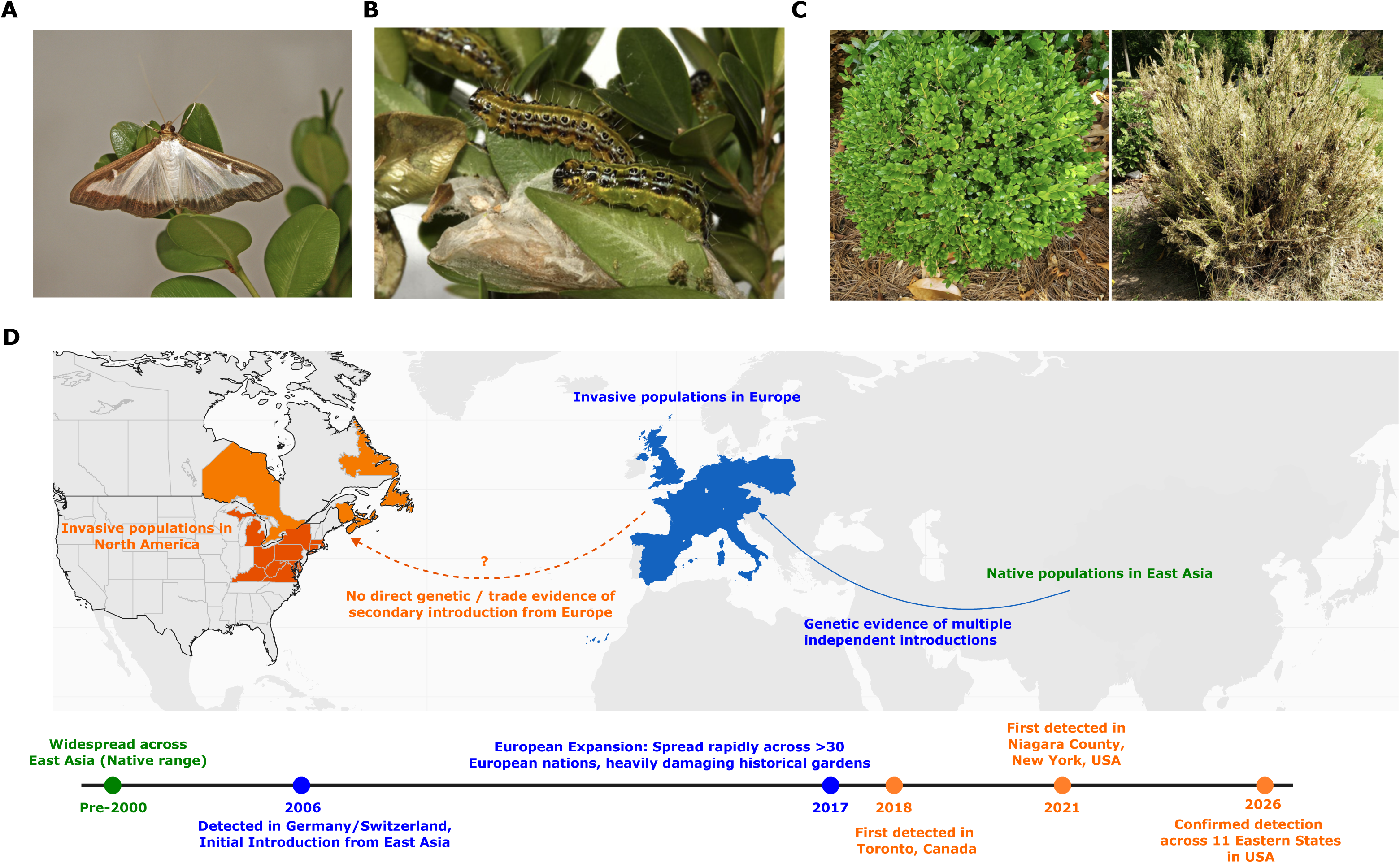
Biology, ecological impact, and global invasion history of *Cydalima perspectalis*. **(A)** Adult box tree moth, an invasive lepidopteran species responsible for widespread damage to ornamental and native boxwood (*Buxus* spp.) trees © USDA. **(B)** *C. perspectalis* larva feeding on boxwood foliage, representing the primary stage responsible for host defoliation and plant damage. **(C)** Impact of *C. perspectalis* infestation on boxwood health. The left panel shows a healthy boxwood tree, whereas the right panel shows a severely damaged boxwood tree following infestation and repeated larval feeding. **(D)** Historical invasion pathway and geographic expansion of *C. perspectalis* from its native range in East Asia to introduced populations in Europe and North America. The map summarizes major invasion events across Europe, Canada, and the United States. The timeline illustrates the progression of global spread and provides the historical framework for investigating genomic signatures of introduction, population differentiation, and demographic change.

The recent and geographically expanding distribution of *C. perspectalis* in North America provides an opportunity to investigate invasion processes while populations are still becoming established. In particular, determining whether newly detected populations originated through a common introduction pathway or through multiple independent introductions is important for understanding the subsequent spread of the invasive species. Human-mediated movement of infested boxwood plants provides a plausible mechanism for both long-distance introduction and secondary spread. However, geographic proximity alone may not accurately reflect invasion history because populations established through separate introductions can occur in neighboring regions, whereas populations separated by large geographic distances may share recent ancestry if connected through plant trade or transportation networks. Understanding invasion pathways is therefore a critical component of predicting and managing biological invasions (Allendorf and Lundquist 2003; Bras et al. 2022). Genome-wide approaches provide powerful tools for reconstructing invasion histories, characterizing population structure and demographic history, identifying genetically distinct introduction lineages, and detecting patterns of admixture and asymmetric allele sharing among introduced populations (Jeffery et al. 2017; McGaughran et al. 2024). Such approaches can complement conventional biosecurity surveillance by providing genetic signatures that can be used to distinguish newly introduced populations from secondary spread of previously established lineages.

Previous genetic studies of *C. perspectalis* in Europe, based primarily on mitochondrial markers (Matošević et al. 2017; Bras et al. 2019a) and microsatellite data (Bras et al. 2022), have suggested introductions from eastern Asia followed by secondary invasions and bridgehead effects, in which established invasive populations have contributed to subsequent introductions into new areas (Bertelsmeier et al. 2018a). Admixture among introduced populations has also been suggested, potentially facilitated by continued movement of *Buxus* plants within Europe (Bras et al. 2022; Seehausen et al. 2024a). These studies demonstrate that invasion histories of *C. perspectalis* can involve multiple introductions, secondary spread, and genetic admixing. A recent ecological modelling study has further indicated that the North American invasion has substantial potential for continued expansion: an updated CLIMEX model predicted that much of North America is climatically suitable for establishment of *C. perspectalis*, with particularly broad areas of suitability across central and eastern North America (Seehausen et al. 2024b). Despite this high potential for further expansion, the genetic processes underlying the early establishment and geographic spread of *C. perspectalis* in North America remain largely unresolved. In particular, it is unknown whether recently detected populations in North America represent expansion from a common introduction source, independent introductions associated with plant trade, or subsequent admixture among introduced populations. Resolving these alternatives is critical for reconstructing invasion pathways and developing effective surveillance and management strategies.

In this study, we used genome-wide single nucleotide polymorphism (SNP) data generated using double-digest restriction-site associated DNA sequencing (ddRAD-seq) to investigate the population genomic history of *C. perspectalis* across its native and introduced ranges. We complemented genome-wide SNP analyses with mitochondrial *COI* gene sequencing and morphometric measurements to provide independent perspectives on geographic differentiation and invasion history. Specifically, we aimed to: (1) characterize patterns of genome-wide genetic diversity and differentiation among native and invasive populations; (2) resolve population structure and ancestry among recently established North American populations and compare their genomic relationships with European and native-range populations; (3) identify patterns of asymmetric allele sharing and demographic history that can distinguish shared ancestry from potentially independent introduction histories; and (4) integrate genomic, mitochondrial, and morphological patterns to identify invasion signatures and evaluate their implications for reconstructing the establishment and spread of *C. perspectalis* in North America. By linking population genomic patterns with known invasion pathways and horticultural trade, we sought to establish a genetic framework that can support surveillance, pathway tracing, and management of this emerging invasive pest.

## Materials and methods

### Sampling

Specimens of *Cydalima perspectalis* were collected between 2024 and 2025 from multiple regions representing (a) recently invaded populations (North America), (b) long-established invasive populations (Europe), and (c) the native range (China) **(Fig. 2A**, **Table 1)**. In the United States, adult moths were sampled from each of the following states: Michigan, Massachusetts, Maryland, Ohio, New York, West Virginia, and Delaware. In addition, pupal specimens were collected from Virginia. Canadian populations were represented by individuals collected from Ontario. Long-established invasive populations in Europe were represented by individuals collected from Switzerland and individuals from the Croatian colony maintained at the Forest Pest Methods Laboratory (FPML), USDA PPQ. To represent the native range, three museum specimens originating from China were included in the genomic study.

**Figure 2.**
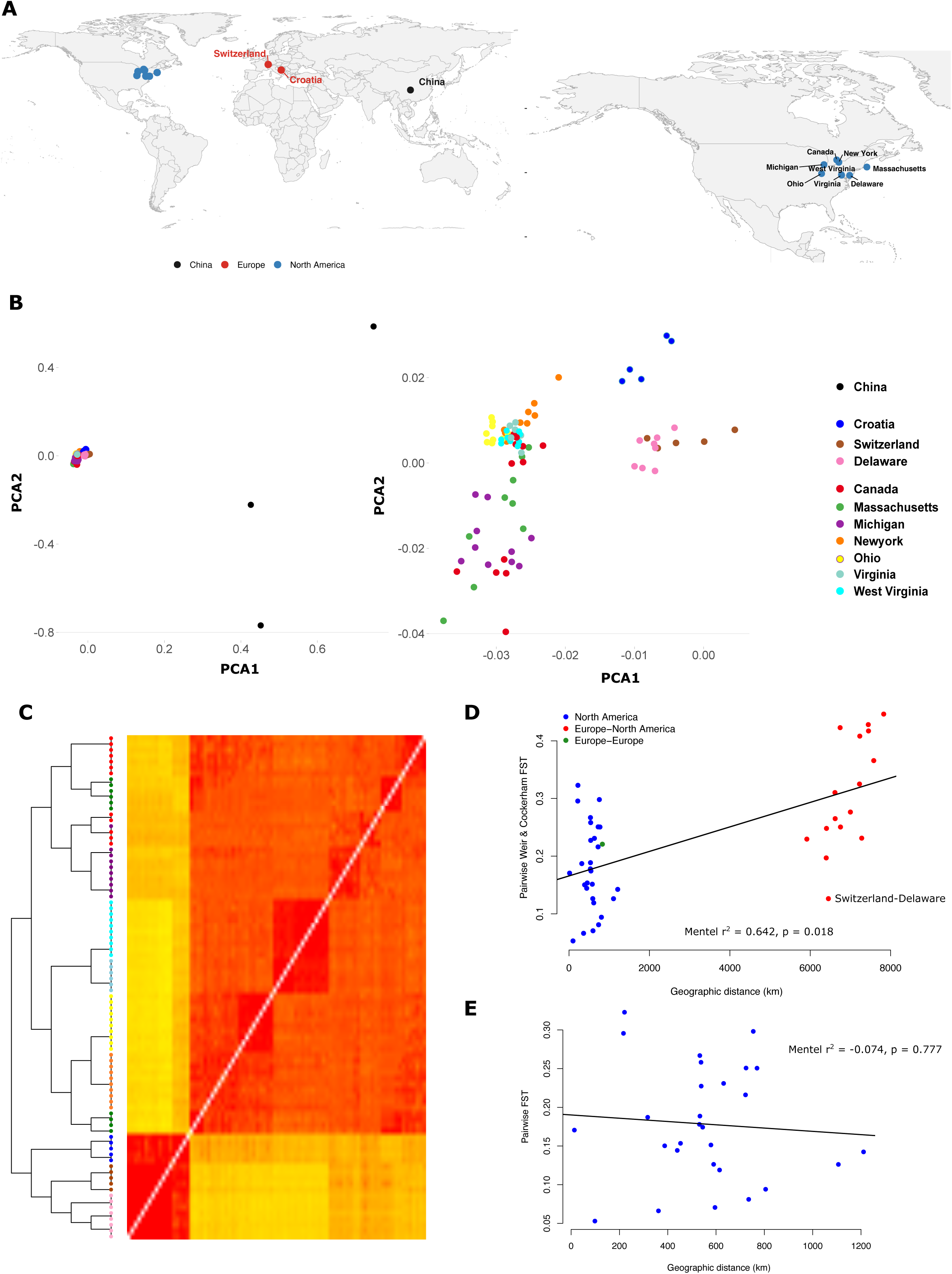
Population genetic structure and geographic patterns of genetic variation in *C. perspectalis*. **(A)** Sampling locations of all populations included in this study. The left panel shows the global distribution of sampled populations, including the native-range population from China and introduced populations from Europe and North America. The right panel shows an expanded view of the North American sampling locations. **(B)** Principal component analysis (PCA) based on 32,712 filtered biallelic SNPs. The left panel includes all sampled populations, illustrating the broad genetic differentiation between the native Chinese population and introduced populations. The right panel excludes the Chinese population to highlight finer-scale genetic relationships among European and North American populations, including the close affinity of the Delaware population with Switzerland. **(C)** Individual co-ancestry matrix inferred using fineRADstructure. Rows and columns represent individuals; red colors indicate greater shared co-ancestry, and yellow represents low shared co-ancestry. The accompanying hierarchical clustering identifies major genetic groups and reveals the close clustering of Delaware individuals with the Swiss population. All populations are coded by the same colors in Fig B and C. **(D)** Isolation-by-distance (IBD) analysis across all sampled populations showing the relationship between pairwise genetic differentiation (F_ST_) and geographic distance. A significant positive Mantel correlation indicates increasing genetic differentiation with geographic distance across the global dataset. **(E)** Isolation-by-distance analysis restricted to North American populations. No significant relationship was detected between geographic and genetic distance, indicating that isolation by distance does not explain genetic structure within the invaded North American range.

**Table 1:** Sampling localities and sample size of *C. perspectalis* populations included in this study.

| Population | mtDNA<br>barcoding<br>sample size | Genomic<br>analysis<br>sample size | Population type | Year first detected | Year sampled |
| --- | --- | --- | --- | --- | --- |
| New York | 16 | 10 | recently invaded<br>(North America) | 2021 (Proxmire, 2022) | 2024 |
| Ohio | 16 | 10 | recently invaded<br>(North America) | 2023 (Boggs et al., | 2024 |
|  |  |  |  | 2025) |  |
| Massachusetts | 20 | 10 | recently invaded<br>(North America) | 2023 (Boggs et al.,<br>2025) | 2024 |
| Michigan | 0 | 10 | recently invaded<br>(North America) | 2022 (Boggs et al.,<br>2025) | 2024 |
| Virginia | 6 | 6 | recently invaded<br>(North America) | 2025 (VDACS, 2025) | 2025 |
| West Virginia | 7 | 10 | recently invaded<br>(North America) | 2025 (WVDA, 2025) | 2025 |
| Maryland | 8 | 0 | recently invaded<br>(North America) | 2025 (MDA, 2025) | 2025 |
| Delaware | 7 | 8 | recently invaded<br>(North America) | 2024 (Boggs et al.,<br>2025) | 2025 |
| Canada | 25 | 12 | recently invaded<br>(North America) | 2018 (Wiesner et al.,<br>2021) | 2024 |
| Switzerland | 0 | 5 | long-established<br>invasive<br>(European) | 2007 (Billen, 2007) | 2024 |
| Croatia | 16 | 5 | long-established<br>invasive<br>(European) | 2012 (Koren & Črne,<br>2012) | 2024 |
| China | 0 | 3 | native | N/A | 2025 |

### Mitochondrial COI barcoding

Genomic DNA was isolated from a leg or antenna using the DNeasy Blood and Tissue Kit (QIAGEN, Alameda, CA). To assess mitochondrial (mtDNA) diversity of the recently invaded North American populations, the 658 bp invertebrate *COI* barcode region was amplified with the universal primer set LCO1490/HCO2198 (Folmer et al. 1994). Each 20 µl PCR contained 9 µl of molecular grade water, 2 µl of 10X PCR buffer lacking MgCl□, 2.8 µl of 25 mM MgCl□, 3.2 µl of 1.25 mM dNTPs, 0.4 µl of each primer (10 pmol/µl), and 0.2 µl of JumpStart Taq DNA Polymerase (Sigma Aldrich, St. Louis, MO; 2.5 units/µl). Two microliters of extracted DNA served as the template. Thermocycling consisted of an initial 2□min denaturation at 94□°C, followed by 40 cycles of 94□°C for 15 s, 52□°C for 30 s, and 72□°C for 1 min, with a final 5□min extension at 72□°C. Negative controls were included to detect potential contamination. PCR products were visualized on 3% agarose gels, purified with ExoSAP IT (Affymetrix, Cleveland, OH), and sequenced on an ABI 3730XL platform (ACGT, Inc., Wheeling, IL). Resulting sequences were manually curated and aligned in Geneious Prime 2024.0.7. To assess haplotype relationships, two mitochondrial genomes (MH602288 and KY865331) that each represented one of the two most prevalent haplotype groups across both native and invaded regions (HTA group and HTB group, respectively) (Gao et al. 2023) were downloaded from GenBank and compared with sequences generated from North American specimens.

### Restriction-site-associated DNA sequencing, alignment, and variant calling

Genomic DNA was extracted from abdominal tissue using the Qiagen DNeasy Blood and Tissue Kit (Qiagen, Cat. No. 69504) following the manufacturer’s animal tissue extraction protocol, including an overnight proteinase-K digestion step. DNA concentration and quality were quantified using a Qubit 4 Fluorometer (Invitrogen, Thermo Fisher Scientific). Double-digest restriction-site-associated DNA sequencing (ddRADseq) libraries were prepared using the EcoRI–MspI restriction enzyme pair following a modified protocol based on (Peterson et al. 2012). Libraries were sequenced on an Illumina Novoseq platform to generate paired-end reads of 2 × 150 bp. Demultiplexed FASTQ files were used for downstream quality filtering, alignment, and variant-calling analyses.

Bioinformatic processing of raw ddRAD-seq reads and genotype calling was performed using STACKS2 v2.68 (Normandeau 2024). Raw reads were demultiplexed, barcode sequences were removed, and low-quality reads were filtered using the ‘process_radtags’ module. Following inspection of per-base quality profiles from the 150 bp paired-end reads, sequences were trimmed to 129 bp prior to downstream analyses. Trimmed paired-end reads were aligned to the publicly available *C. perspectalis* reference genome (Broad et al. 2024) using BWA-MEM v0.7.17 (Li and Durbin 2009). Resulting BAM files were filtered to retain reads with a minimum mapping quality score of 10, then sorted and indexed using SAMtools v1.16.1 (Li et al. 2009). SNP discovery and genotype calling were conducted using the ‘gstacks’ and ‘populations’ modules implemented in STACKS2 v2.68 (Normandeau 2024). Variant filtering was subsequently performed using STACKS2 and VCFtools (Danecek et al. 2011). SNPs with greater than 30% missing data per site, a minor allele count less than 4, or a variant quality score below 20 were excluded. The final dataset consisted of 32,712 high-quality biallelic SNPs from 89 individuals and was used for downstream population genomic analyses.

### Principal component analysis (PCA)

The PCA was conducted from the final filtered SNP dataset in PLINK v1.9 (Chang et al. 2015) using the --pca function, and the resulting principal component coordinates were visualized in R (R Core Team 2024) using ggplot2. To minimize the effects of linked SNPs, SNPs were pruned in PLINK using a sliding-window approach with a window size of 50 SNPs, a step size of 10 SNPs, and a linkage disequilibrium (r²) threshold of 0.1. After pruning, 2,544 SNPs were retained for PCA. To evaluate genomic relationships across the entire sampled range, an initial PCA included all individuals, including the three native specimens. Because the native samples formed a highly divergent cluster relative to the introduced populations, which appeared tightly clustered in the PCA, a second PCA was performed after excluding the native samples to improve resolution of population structure within the invasive European and North American populations. Principal components were plotted using the first two axes (PC1 and PC2), which explained the largest proportion of genetic variation.

Population structure and ancestry proportions were further inferred using ADMIXTURE v1.3.0 (Alexander et al. 2009) based on the 32,712 genome-wide SNPs. The optimal number of genetic clusters (K) was evaluated using the cross-validation procedure implemented in ADMIXTURE. Cross-validation error was assessed across multiple K values, with the lowest error observed at K = 4, followed by K = 2. The selected models were subsequently used to estimate ancestry proportions for each individual, and admixture proportions were visualized across all sampled populations.

### fineRADstructure analysis

To investigate recent shared ancestry and fine-scale population structure, we analyzed the filtered 32,712 genome-wide SNP dataset using fineRADstructure v0.3.2 (Malinsky et al. 2016). The co-ancestry matrix was first estimated using RADpainter, which infers nearest-neighbor haplotype sharing among individuals based on RAD loci. The resulting co-ancestry matrix was then clustered using the fineRADstructure Markov chain Monte Carlo (MCMC) algorithm. The MCMC analysis was run with 100,000 burn-in iterations followed by 100,000 sampling iterations, with samples recorded every 1,000 iterations. Population relationships were summarized using the maximum-clade-credibility tree generated by fineRADstructure. The co-ancestry heatmap and hierarchical clustering tree were visualized using the R scripts provided with the fineRADstructure package.

### Isolation-by-distance analysis

To evaluate whether genetic differentiation increased with geographic distance, we tested for isolation by distance (IBD) using Mantel tests. Pairwise genetic differentiation among populations was quantified with F_ST_ values estimated from the filtered SNP dataset with VCFtool (Danecek et al. 2011) using the Weir and Cockerham estimator. Geographic distances between sampling locations were calculated as great-circle (geodesic) distances based on latitude and longitude coordinates using the geosphere package in R (R Core Team 2024).

The relationship between pairwise genetic distance (F_ST_) and geographic distance was assessed using Mantel tests implemented in the vegan package in R with 9,999 permutations to evaluate statistical significance. Two analyses were performed: (i) an analysis including all sampled European and North American populations to evaluate isolation by distance across the invaded range, and (ii) a second analysis restricted to North American populations to determine whether geographic distance explained genetic differentiation within the invaded North American range. Scatterplots showing the relationship between geographic distance and pairwise F_ST_ were generated using ggplot2, with linear regression lines added for visualization.

### D-statistic (Dsuite) analysis

The D-statistic provides a test of asymmetric allele sharing among populations and can be used to identify patterns consistent with gene flow, admixture, or shared ancestry that are not explained by a simple bifurcating population history (Green et al. 2010). In the context of invasion biology, these patterns can help distinguish a single introduction followed by geographic expansion from more complex invasion histories involving multiple introductions, secondary contact, or genetic admixing among introduced populations (Sillo et al. 2021). Although D-statistics do not directly estimate migration rates, significant excess allele sharing can provide evidence for historical or ongoing genetic connectivity among populations.

Patterns of allele sharing among invasive populations were evaluated using the ABBA–BABA framework implemented in Dsuite (Malinsky et al. 2021). Genome-wide D-statistics were calculated using the Dtrios function from the filtered SNP dataset to test all possible population quartets. Because the three native samples contained a high proportion of missing data and reduced the number of informative SNPs, they were excluded from these analyses. European populations from Croatia and Switzerland were used independently as alternative outgroup (P4) populations to evaluate the robustness of inferred allele-sharing relationships. Statistical significance was assessed using block-jackknife Z-scores, which account for linkage among neighboring SNPs by recalculating D-statistics after sequentially removing contiguous genomic blocks. Z-scores generated from the block-jackknife analysis were used to assess significance, and quartets with |Z-score| ≥ 3 were considered to show significant asymmetric allele sharing. Positive D-statistics (excess ABBA patterns) indicate excess allele sharing between P2 and P3, whereas negative D-statistics (excess BABA patterns) indicate excess allele sharing between P1 and P3.

To summarize recurrent patterns of asymmetric allele sharing among invasive populations, only significant Dsuite tests (|Z-score| ≥ 3) were retained. Because significant D-statistics indicate excess derived allele sharing between the P2 and P3 populations relative to P1, only P2–P3 population pairs were considered when constructing the summary bar plot. Each significant quartet contributed one count to the corresponding P2–P3 population pair, irrespective of the outgroup (P4) population (Croatia or Switzerland). Counts were then summed across all significant quartets to quantify the frequency with which each pair of populations exhibited significant excess allele sharing.

### Estimates of population genetic diversity and inbreeding

Within-population genetic diversity was quantified using nucleotide diversity (π), observed heterozygosity (Ho), and the inbreeding coefficient (F). Population-level nucleotide diversity (π) was estimated from the filtered SNP dataset using VCFtools v0.1.16 (Danecek et al. 2011) with the --site-pi function, and mean nucleotide diversity for each population was calculated by averaging estimates across all polymorphic sites. Observed heterozygosity (Ho) and individual inbreeding coefficients (F) were calculated using PLINK v1.9 (Chang et al. 2015). Observed heterozygosity was estimated for each individual from genome-wide SNP genotypes, and the inbreeding coefficient was calculated using the method-of-moments estimator implemented in PLINK (--het), where positive values indicate heterozygosity deficiency relative to Hardy–Weinberg expectations and negative values indicate excess heterozygosity. Population-level summaries of nucleotide diversity were visualized as Cleveland dot plots, whereas the distributions of observed heterozygosity and inbreeding coefficients were summarized using boxplots generated in R with ggplot2.

### Estimates of effective population size and inference of demographic history

Historical changes in effective population size (Ne) were inferred using GONE (Santiago et al. 2020), which estimates recent demographic history from patterns of linkage disequilibrium among genome-wide SNPs. GONE estimates temporal changes in effective population size without requiring phased genotypes by modeling the decay of linkage disequilibrium across recombination distances. Separate analyses were performed for each population using the filtered SNP dataset. Because GONE requires physical marker positions, SNP coordinates from the *C. perspectalis* reference genome (Broad et al. 2024) were retained in the input files. Analyses were conducted using default parameters, and historical effective population size trajectories were reconstructed across recent generations. Estimated effective population size trajectories were visualized in R using ggplot2.

### Specimen preparation and morphometric analysis

Adult *C. perspectalis* specimens were prepared for morphometric analysis following standard entomological preservation and mounting procedures previously used for *C. perspectalis* (Altun et al. 2026). Morphological measurements were obtained from dried adult specimens using standardized procedures. Six morphological traits were measured: head length (distance between the anterior and posterior margins of the head capsule), head width (maximum lateral width of the head capsule), body length (distance from the anterior margin of the head to the posterior tip of the abdomen), forewing length (wing base to apex), hindwing length (wing base to apex), and wingspan (maximum distance between the apices of fully expanded forewings). All measurements were recorded to the nearest 0.01 mm using digital calipers.

Morphological variation among populations was assessed using one-way analysis of variance (ANOVA) for each trait. Assumptions of normality and homogeneity of variance were evaluated using the Shapiro–Wilk and Levene’s tests, respectively. When homogeneity of variance was violated, Welch’s ANOVA was used to confirm statistical significance, and when residuals deviated from normality, results were additionally verified using the Kruskal–Wallis test (Liu 2015). Pairwise differences among populations were evaluated using Tukey’s honestly significant difference (HSD) test with adjustment for multiple comparisons. Relationships among morphological traits were examined using Pearson correlation coefficients. Principal component analysis (PCA) was performed on standardized trait measurements to summarize multivariate patterns of morphological variation among populations. All statistical analyses and visualizations were conducted in R using the packages ggplot2, ggcorrplot, and FactoMineR/factoextra.

## Results

### Mitochondrial DNA diversity reveals a predominant North American *COI* haplotype

Mitochondrial *COI* sequences were obtained from 105 *C. perspectalis* specimens representing seven US states and Ontario, Canada, together with 16 specimens from the Croatian FPML laboratory colony. Mitochondrial haplotype analysis revealed two major COI haplotype groups, HTA and HTB. The HTA haplotype group was detected only in the Croatian colony and Delaware specimens. In contrast, all remaining sampled US populations and the Canadian population shared an identical COI barcode belonging to the HTB haplotype group.

### Reduced genetic diversity and population genetic structure following invasion

We further expanded mtDNA to genome-wide analysis and analyzed populations of the *C. perspectalis* sampled across its invaded range in North America and Europe and compared them with native populations from China **(Fig. 2A)**. In total, genomic analyses (using ddRAD sequencing) were performed using 89 individuals (Table 1) after quality filtering and processing, resulting in a final dataset of 32,712 high-quality biallelic genome-wide SNPs. These SNPs were used consistently across downstream population genomic analyses, including population structure analyses, allele-sharing tests, genetic differentiation estimates, and demographic inference. The PCA revealed a clear contrast between the native-range population and populations sampled from the introduced ranges across Europe and North America **(Fig. 2B, left panel)**. Native range individuals showed a broader distribution across PCA space, indicating greater genetic variation within the native population. In contrast, European and North American invasive populations formed a comparatively compact cluster, suggesting reduced genome-wide variation and high genetic similarity among introduced populations.

### Distinct introduction histories among invasive populations

To further resolve genetic relationships among introduced populations, we performed a separate PCA excluding the genetically divergent native-range population **(Fig. 2B, right panel)**. This analysis revealed finer-scale differentiation among European and North American populations and highlighted distinct patterns of genetic affinity within the introduced range. European populations (Croatia and Switzerland) formed a separate cluster from most North American populations, indicating genetic differentiation between European and North American lineages. North American populations did not form strongly defined location-specific clusters. Instead, individuals from most North American locations showed a broad distribution across PCA space, with substantial overlap among populations.

The Delaware population, however, showed a contrasting pattern, forming a relatively compact cluster that was positioned closer to the European populations than to most other North American populations. Among European populations, Delaware individuals showed the strongest affinity toward the Switzerland population in PCA space. To resolve fine-scale relationships among introduced populations, we further analyzed individual-level genetic relatedness using fineRADstructure **(Fig. 2C)**. European populations formed a distinct co-ancestry group, consistent with their separation in PCA analyses. Notably, individuals from Delaware were placed within the same hierarchical cluster as the Swiss population, showing elevated co-ancestry with Switzerland compared with other North American populations. In contrast, individuals from most other North American populations did not form strongly separated state-specific clusters. Instead, they showed broader patterns of shared ancestry and mixed placement within the North American cluster, consistent with the diffuse structure observed in PCA. Together, these results suggest that Delaware has a different genetic history from other North American populations and may retain stronger ancestry from a European source population.

### Isolation by distance is evident globally but absent within the North American invasion

To evaluate the relationship between geographic distance and genetic differentiation, we performed Mantel tests using pairwise geographic distances and pairwise F_ST_ values. Across all sampled populations, including Europe and North America, genetic differentiation increased significantly with geographic distance (Mantel r^2^ = 0.642, p = 0.018; **Fig. 2D**). This positive correlation indicates that populations separated by greater geographic distances also tend to be more genetically differentiated, reflecting broad-scale geographic structuring across the Box Tree Moth invasive ranges. In contrast, when the analysis was restricted to North American populations, no significant relationship was detected between geographic and genetic distance (Mantel r^2^ = −0.074, p = 0.777; **Fig. 2E**). Genetic differentiation among North American populations was therefore independent of geographic proximity, indicating that isolation by distance does not explain population genetic structure within the invaded range.

The absence of isolation by distance within North America suggests that the observed genetic structure is unlikely to have arisen through gradual spatial expansion following a single introduction. Instead, the result is more consistent with multiple introduction events and/or long-distance human-mediated dispersal. An exception to the general North American pattern was the Delaware population, which consistently showed greater genetic affinity to the Swiss population in PCA, fineRADstructure, and IBD analyses than to other North American populations, suggesting a distinct introduction history. However, because the sampling design was geographically unbalanced, with substantially greater representation of North American populations than European reference populations, the IBD analysis should be interpreted cautiously. In particular, the predominance of North American samples may disproportionately influence the relationship between geographic and genetic distances and limit the extent to which the analysis can be generalized across the full native and introduced ranges. The IBD analysis was therefore used primarily as a complementary assessment of geographic structuring rather than as an independent test of population structure.

### Genome-wide ancestry reveals heterogeneous genetic structure among invasive populations

To investigate population ancestry and genetic structure within the invasive range, we performed model-based clustering using ADMIXTURE, excluding the three genetically divergent native samples. Cross-validation analysis identified K = 4 (CV error = 0.383) as the best-supported model, followed closely by K = 2 (CV error = 0.397). At K = 2, invasive populations were divided into two major genetic clusters **(Fig. 3A, upper panel)**. One cluster comprised European and Delaware populations from North America. The second cluster comprised the remaining North American populations, indicating that Delaware shares substantially greater ancestry with European populations than other invasive North American populations.

**Figure 3:**
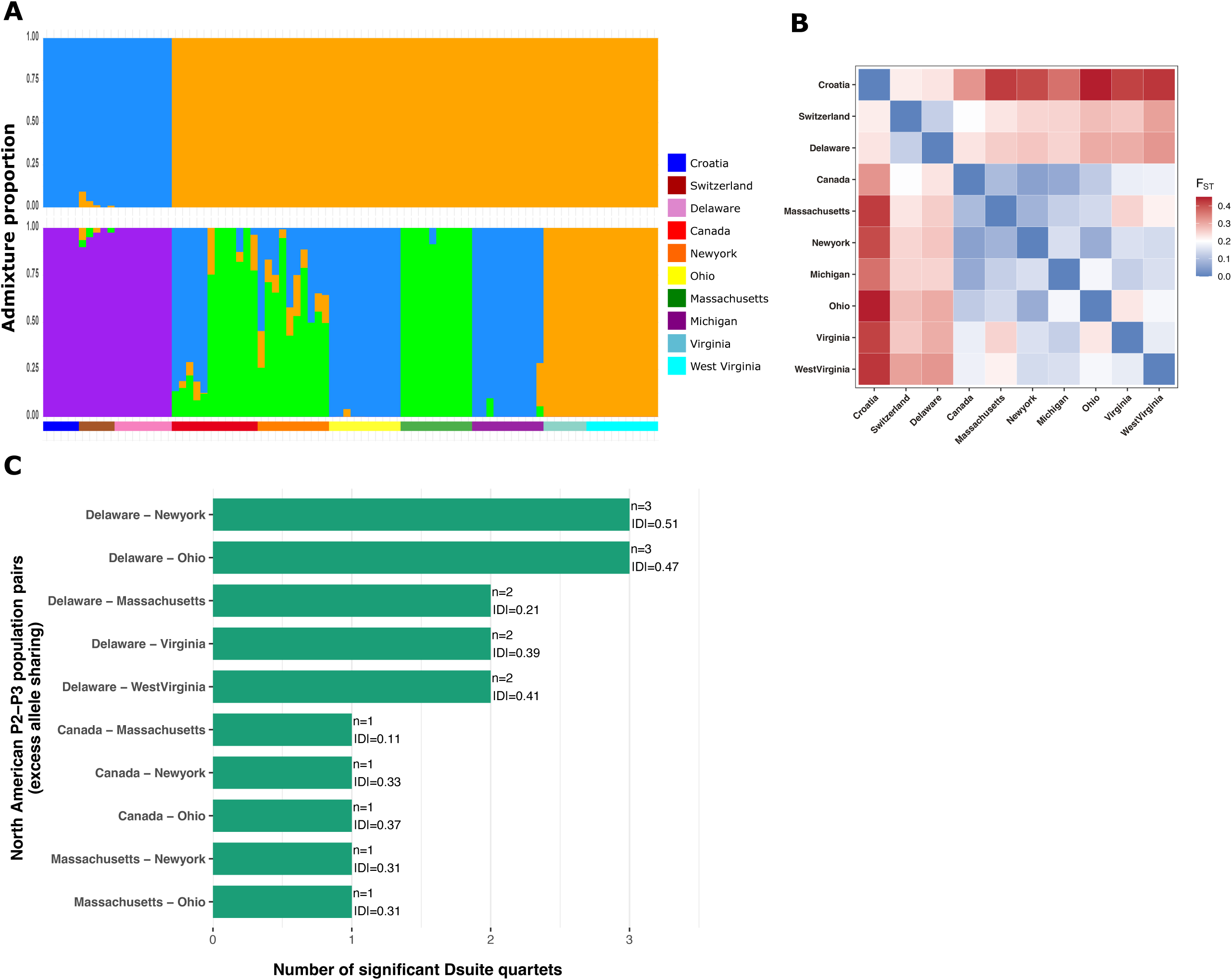
Genomic ancestry, population differentiation, and asymmetric allele sharing among invasive *C. perspectalis* populations. **(A)** ADMIXTURE analysis of invasive populations (native population from China excluded) showing individual ancestry proportions at K = 2 (upper panel) and K = 4 (lower panel), corresponding to the lowest cross-validation error values. Each vertical bar represents one individual, and colors indicate the inferred proportion of ancestry assigned to each genetic cluster. Populations are grouped by sampling locality. **(B)** Heatmap of pairwise genetic differentiation (mean F_ST_) among invasive populations. Warmer colors indicate greater genetic differentiation. Delaware exhibits lower differentiation from the European populations, particularly Switzerland, than from most other North American populations. **(C)** Frequency of significant excess allele-sharing relationships among North American population pairs identified using Dsuite analyses. Bars represent the number of significant quartets (|Z| ≥ 3) in which each North American population pair occurred as the P2– P3 pair, indicating repeated excess allele sharing relative to alternative North American populations (P1) and using either Croatia or Switzerland populations as outgroup (P4). Values adjacent to each bar indicate the number of supporting quartets and the mean absolute D-statistic (|D|) across those significant tests. Frequent involvement of Delaware (5 out of 10) in significant P2–P3 pairings identifies it as a recurrent participant in asymmetric allele-sharing relationships with multiple North American populations, consistent with a more complex introduction history than observed for most other invasive North American populations.

At K = 4 **(Fig. 3A, lower panel)**, finer-scale population genetic structure became evident. European populations and Delaware continued to share a common ancestry component that remained distinct from the rest of North America. Within North America, three largely non-admixed ancestry groups were identified: (i) Ohio and Michigan, (ii) Massachusetts, and (iii) Virginia and West Virginia. In contrast, individuals from Canada and New York exhibited admixed ancestry, indicating genetic mixing among North American lineages following introduction.

### Pairwise genetic differentiation supports heterogeneous population genetic structure within the invaded range

Pairwise F_ST_ estimates revealed substantial variation in genetic differentiation among introduced populations, consistent with the ancestry patterns inferred from ADMIXTURE analysis **(Fig. 3B)**. The lowest levels of differentiation were observed among several North American populations, including Canada–New York ( F_ST_ = 0.053), Canada– Michigan (0.066), New York–Ohio (0.071), Massachusetts–New York (0.081), and Canada– Massachusetts (0.094), indicating relatively high genetic similarity among these populations. In contrast, the Delaware population showed a distinct pattern. Despite its geographic location within North America, Delaware exhibited its lowest genetic differentiation with the Switzerland population (F_ST_ = 0.126), whereas differentiation between Delaware and other North American populations ranged from 0.231 to 0.323. This pattern closely mirrors the PCA, fineRADstructure, IBD, and ADMIXTURE analyses, which also placed Delaware in close association with the Swiss populations.

Among European populations, Croatia exhibited the greatest genetic differentiation from most North American populations (F_ST_ = 0.325–0.446), while showing comparatively lower differentiation from Switzerland (0.221) and Delaware (0.231). Within North America, moderate differentiation was observed among several regional populations, supporting the presence of multiple genetic groups within the invaded range.

### Repeated asymmetric allele sharing among North American populations

To evaluate patterns of allele sharing among invasive populations, we performed ABBA/BABA analyses using Dsuite and tested all possible population quartets. European populations (Croatia and Switzerland) were used independently as alternative P4 (outgroup) populations to evaluate the robustness of inferred relationships. The highest Z-score was observed for the quartet Michigan– Ohio–Delaware (D = 0.697, Z = 13.33), followed by Michigan–NewYork–Delaware (D = 0.680, Z = 10.97) **(Supplementary Table 1)**.

To examine the recurrent patterns of asymmetric allele sharing within the introduced range, we summarized the frequency of significant Dsuite quartets involving each North American P2–P3 population pair among the significant ABBA/BABA tests. Significant excess allele-sharing signals were not evenly distributed among population pairs but were concentrated among a subset of recurrent relationships. Notably, Delaware was involved in five of the ten most frequently observed significant P2–P3 relationships, including repeated allele-sharing associations with Ohio, New York, Massachusetts, Virginia, and West Virginia **(Fig. 3C)**. This recurrent involvement of Delaware across multiple independent quartet tests suggests that this population shares ancestry components with multiple invasive lineages rather than reflecting a single, isolated introduction history. Other frequently observed relationships included allele-sharing signals among northeastern populations (e.g., Canada, Massachusetts, New York, and Ohio), further indicating that North American populations do not represent a single genetically uniform invasion lineage.

### Genetic diversity and demographic history reveal contrasting invasion dynamics among North American populations

Nucleotide diversity (π) varied substantially among invasive populations, revealing pronounced differences in the amount of standing genetic variation retained following introduction **(Fig. 4A)**. Among the invasive populations, the two European populations exhibited contrasting levels of diversity, with Switzerland showing the highest nucleotide diversity overall (π = 6.77 × 10□□), followed by Croatia (π = 4.57 × 10□□). Within North America, Delaware maintained the highest nucleotide diversity (π = 5.57 × 10□□), exceeding all other introduced populations and approaching the level observed in Switzerland. Intermediate levels of diversity were observed in New York (4.53 × 10□□), Ohio (4.51 × 10□□), Virginia (4.44 × 10□□), West Virginia (4.01 × 10□□), and Michigan (3.34 × 10□□), whereas Massachusetts (1.72 × 10□□) and Canada (1.68 × 10□□) exhibited the lowest nucleotide diversity among all invasive populations.

**Figure 4.**
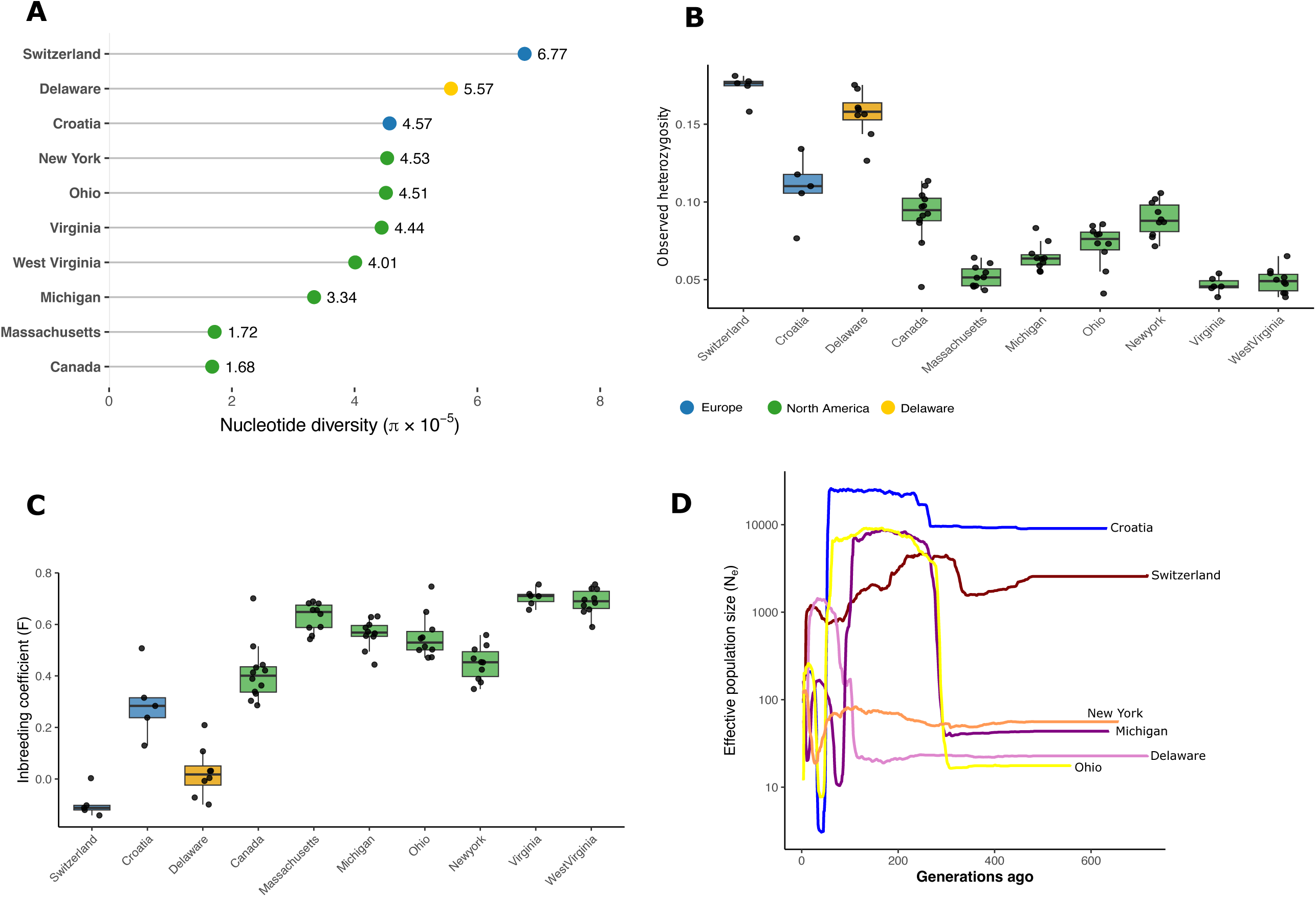
Genetic diversity and demographic history of invasive *Cydalima perspectalis* populations. **(A)** Population-level nucleotide diversity (π) estimated from genome-wide ddRADseq SNP variation. Populations are ranked according to mean π values. Delaware exhibited the highest nucleotide diversity among North American populations, approaching levels observed in European populations. **(B)** Distribution of individual observed heterozygosity (Ho) values across sampled populations. Boxplots represent individual-level genome-wide heterozygosity estimates, with center lines indicating medians, boxes representing the interquartile range, and whiskers showing the distribution of values excluding outliers. Black dots represent observed heterozygosity of individual samples. Delaware individuals showed elevated heterozygosity relative to most North American populations. The same color coding is used for Fig. A, B**, and C. (C)** Distribution of individual inbreeding coefficients (F) among populations. Boxplots represent genome-wide individual estimates of inbreeding. Black dots represent observed heterozygosity of individual samples. Delaware exhibited consistently low inbreeding values, whereas several North American populations showed higher heterozygosity deficits. **(D)** Historical effective population size (Ne) trajectories inferred using GONE from linkage disequilibrium decay patterns. Lines represent population-specific estimates of effective population size through time, with more recent generations shown toward the left side of each plot. Representative North American populations are shown to highlight contrasting demographic trajectories; complete GONE results for all populations are provided in Supplementary Figure X. European populations showed contrasting demographic histories, with the laboratory-maintained Croatia population exhibiting a recent reduction consistent with colony establishment and maintenance, whereas Switzerland retained higher effective population size estimates. Among North American populations, Michigan and Ohio displayed similar trajectories characterized by population expansion beginning approximately 250 generations ago followed by recent decline, while Delaware exhibited a contrasting pattern of recent effective population size increase during the period when Midwestern populations declined.

Observed heterozygosity showed a pattern similar to nucleotide diversity **(Fig. 4B)**. Switzerland exhibited the highest heterozygosity (Ho = 0.174), followed by Delaware (Ho = 0.156), whereas all remaining North American populations exhibited substantially lower heterozygosity (0.047– 0.092). Croatia showed intermediate heterozygosity (Ho = 0.109). Within North America, Massachusetts (Ho = 0.052), West Virginia (Ho = 0.049), and Virginia (Ho = 0.047) exhibited the lowest heterozygosity, consistent with reduced genome-wide genetic variation following introduction.

Estimates of the inbreeding coefficient further distinguished Delaware from the remaining invasive populations **(Fig. 4C)**. Switzerland showed an inbreeding coefficient close to zero, expected for a large population size with greater standing genetic variation, whereas Delaware exhibited the lowest positive inbreeding coefficient among invasive North American populations. In contrast, all other North American populations showed substantially higher inbreeding coefficients. Particularly high inbreeding coefficients were observed in Virginia, West Virginia (0.688), and Massachusetts, indicating pronounced reductions in heterozygosity relative to Hardy–Weinberg expectations. Together with nucleotide diversity estimates, these results indicate that Delaware has retained substantially greater genetic variation than other North American populations.

Furthermore, we reconstructed recent changes in effective population size using GONE, based on patterns of linkage disequilibrium decay from genome-wide ddRAD-seq SNP data. Population-specific trajectories revealed substantial variation in recent demographic histories among the sampled populations **(Fig. 4D)**. The two European populations showed contrasting patterns. The Croatian population, which represents a laboratory-maintained colony, exhibited a relatively large historical effective population size followed by a pronounced decline toward the most recent generations. This recent reduction is likely associated with the establishment and subsequent maintenance of the laboratory colony from a limited breeding population. In contrast, the Swiss population, representing a wild population, maintained comparatively higher effective population size estimates across much of the reconstructed period, although a decline was also evident toward the most recent generations. These contrasting trajectories suggest differences in recent demographic history between the laboratory-maintained and wild European populations. The GONE trajectories for the North American populations were more difficult to interpret in the context of the invasion history. Because *C. perspectalis* was first detected in North America only in 2018, the period since introduction is short relative to the temporal scale over which linkage-disequilibrium-based methods reconstruct demographic changes. Consequently, the trajectories observed for North American populations may largely reflect demographic histories of their source populations prior to introduction rather than demographic changes occurring after establishment in North America.

### Morphological divergence parallels genomic differentiation across invasive populations

To evaluate phenotypic variation among invasive populations, we measured six morphological traits, including head length, head width, body length, forewing length, hindwing length, and wingspan, in 39 *C. perspectalis* individuals collected from seven locations (Switzerland (SW), n = 6, Croatia (CR), n = 6; Canada (CA), n = 4; Massachusetts (MA), n = 5; Michigan (MI), n = 6; New York (NY), n = 6; and Ohio (OH), n = 6;).

Morphological variation differed significantly among sampling locations for five of the six measured traits **(Fig. 5A-F**, **Table 2)**. Wingspan exhibited the strongest geographic differentiation (ANOVA: *F*□,□□= 7.77, *P* < 0.001, η² = 0.59), followed by head width (*F*□,□□= 6.26, *P* < 0.001, η² = 0.54), forewing length (*F*□,□□= 5.75, *P* < 0.001, η² = 0.52), hindwing length (*F*□,□□= 5.08, *P* < 0.001, η² = 0.49), and head length (*F*□,□□= 2.49, *P* = 0.043, η² = 0.32). Body length did not differ significantly among locations (*F*□,□□= 1.42, *P* = 0.236).

**Figure 5.**
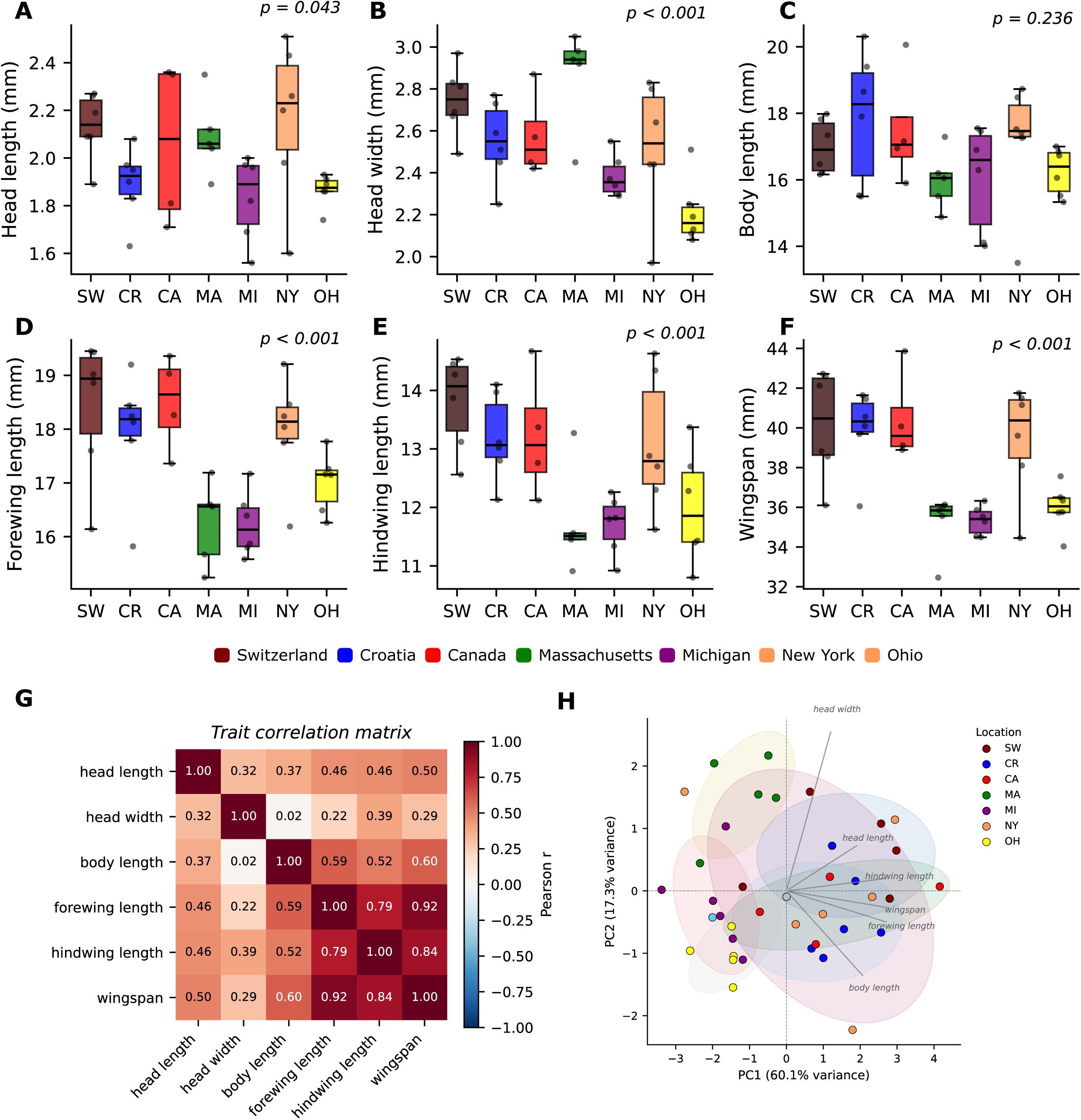
Morphological variation and multivariate differentiation of *C. perspectalis* across seven sampled populations. (A–F) Boxplots of six morphological traits measured from adult moths collected in Canada (CA), Croatia (CR), Massachusetts (MA), Michigan (MI), New York (NY), Ohio (OH), and Switzerland (SW): **(A)** head length, **(B)** head width, **(C)** body length, **(D)** forewing length, **(E)** hindwing length, and **(F)** wingspan. Boxes indicate the median and interquartile range (IQR), whiskers extend to the most extreme observations within 1.5 × IQR, and gray points represent individual measurements (n = 4–6 individuals per population). p-values correspond to one-way ANOVA testing for differences among populations. **(G)** Pearson correlation matrix showing pairwise correlations among all six morphological traits. Correlation coefficients (r) are shown within each cell, with colors representing the direction and strength of the correlation from negative (blue) to positive (red). **(H)** Principal component analysis (PCA) of standardized morphological traits. Points represent individual moths colored by population, and ellipses indicate the 95% confidence interval around each population centroid. Gray arrows represent trait loadings on the first two principal components, and axis labels indicate the percentage of total variance explained by PC1 and PC2.

**Table 2.**
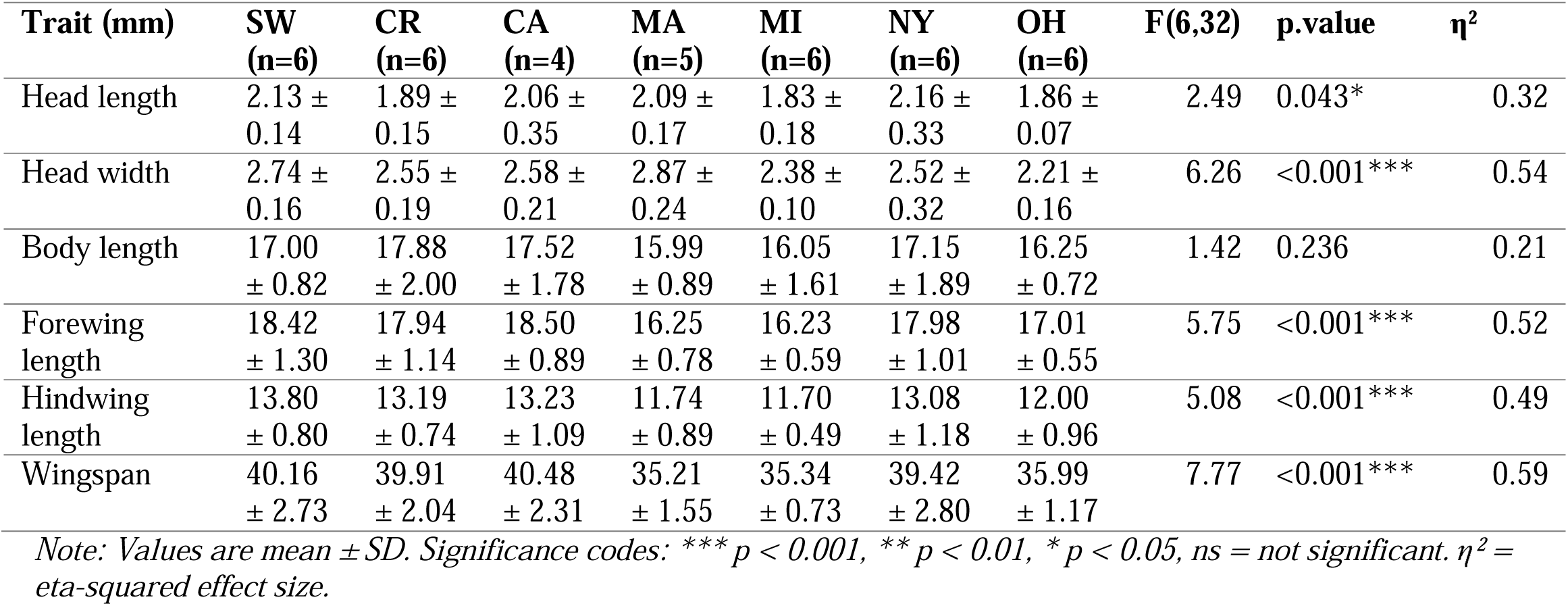
Mean ± SD of morphological traits of *C. perspectalis* by location, with one-way ANOVA results.

| Trait (mm) | SW<br>(n=6) | CR<br>(n=6) | CA<br>(n=4) | MA<br>(n=5) | MI<br>(n=6) | NY<br>(n=6) | OH<br>(n=6) | F(6,32) | p.value | $\eta^2$ |
| --- | --- | --- | --- | --- | --- | --- | --- | --- | --- | --- |
| Head length | 2.13 ± 0.14 | 1.89 ± 0.15 | 2.06 ± 0.35 | 2.09 ± 0.17 | 1.83 ± 0.18 | 2.16 ± 0.33 | 1.86 ± 0.07 | 2.49 | 0.043* | 0.32 |
| Head width | 2.74 ± 0.16 | 2.55 ± 0.19 | 2.58 ± 0.21 | 2.87 ± 0.24 | 2.38 ± 0.10 | 2.52 ± 0.32 | 2.21 ± 0.16 | 6.26 | <0.001*** | 0.54 |
| Body length | 17.00 ± 0.82 | 17.88 ± 2.00 | 17.52 ± 1.78 | 15.99 ± 0.89 | 16.05 ± 1.61 | 17.15 ± 1.89 | 16.25 ± 0.72 | 1.42 | 0.236 | 0.21 |
| Forewing length | 18.42 ± 1.30 | 17.94 ± 1.14 | 18.50 ± 0.89 | 16.25 ± 0.78 | 16.23 ± 0.59 | 17.98 ± 1.01 | 17.01 ± 0.55 | 5.75 | <0.001*** | 0.52 |
| Hindwing length | 13.80 ± 0.80 | 13.19 ± 0.74 | 13.23 ± 1.09 | 11.74 ± 0.89 | 11.70 ± 0.49 | 13.08 ± 1.18 | 12.00 ± 0.96 | 5.08 | <0.001*** | 0.49 |
| Wingspan | 40.16 ± 2.73 | 39.91 ± 2.04 | 40.48 ± 2.31 | 35.21 ± 1.55 | 35.34 ± 0.73 | 39.42 ± 2.80 | 35.99 ± 1.17 | 7.77 | <0.001*** | 0.59 |

Note: Values are mean ± SD. Significance codes: \*\*\* $p < 0.001$ , \*\* $p < 0.01$ , \* $p < 0.05$ , ns = not significant. $\eta^2$ = eta-squared effect size.

Because Levene’s test indicated unequal variances for head length (*P* = 0.035), we confirmed this result using Welch’s ANOVA, which likewise detected significant differences among populations (*F*□,□□□.□= 4.64, *P* < 0.001). Similarly, forewing length showed non-normal residuals (Shapiro–Wilk test, *P* = 0.020), but a non-parametric Kruskal–Wallis test confirmed significant population differences (*H* = 19.39, *P* = 0.004). All remaining traits satisfied assumptions of normality and homogeneity of variance. Post hoc Tukey’s HSD comparisons revealed a consistent geographic pattern in body size. Individuals from Switzerland, Croatia, Canada, and New York generally possessed larger wing dimensions than individuals from Massachusetts, Michigan, and Ohio **(Fig. 5A-F)**. Wingspan differed significantly between Canada and Croatia relative to Massachusetts, Michigan, and Ohio (all adjusted *P* < 0.05), while Switzerland exhibited significantly greater wingspan than both Massachusetts and Michigan (adjusted *P* < 0.01). Forewing and hindwing length followed similar patterns. Although the overall ANOVA for head length was significant, no pairwise comparison remained significant following multiple-test correction.

All six morphological traits were positively correlated **(Fig. 5G)**. The strongest relationships occurred among wing and body-size traits, including forewing length and wingspan (r = 0.92), hindwing length and wingspan (r = 0.84), and forewing length and hindwing length (r = 0.79). In contrast, head width was relatively independent of the other traits and showed the weakest correlation with body length (r = 0.02).

Principal component analysis of standardized morphological traits revealed that the first principal component (PC1) explained 60.1% of the total morphological variation and loaded positively on all six traits, representing a general body-size axis **(Fig. 5H)**. The second principal component (PC2) explained an additional 17.3% of the variation and was driven primarily by head width. Individuals from Massachusetts, Michigan, and Ohio clustered toward negative PC1 values, whereas individuals from Switzerland, Croatia, and Canada clustered toward positive PC1 values, with New York occupying an intermediate position. This multivariate pattern closely mirrored the univariate analyses and indicates that geographic differences among populations are largely explained by overall variation in body and wing size rather than independent divergence of individual morphological traits.

Because the morphological dataset was relatively small and unevenly distributed among locations (n = 4–6 individuals per location), the observed patterns should be interpreted cautiously. Nevertheless, the significant effects detected for five of the six measured traits, the consistency of the geographic pattern across multiple wing and head traits, and the concordance between the univariate analyses and PCA indicate that the observed morphological differentiation is unlikely to be attributable to a single trait or comparison. These results provide evidence of geographic variation in morphology, while larger and more balanced samples will be necessary to determine the extent and stability of these differences across the broader North American invasion.

## Discussion

Minimizing the impacts of invasive alien species (IAS) requires a clear understanding of invasion pathways, introduction history, and the evolutionary processes shaping newly established populations (Pyšek et al. 2020). Such information is particularly valuable during the early stages of an invasion, when targeted management interventions may be most effective in limiting establishment and further spread (Liebhold and Tobin 2008). In this study, we investigated the early-phase invasion of the box tree moth *(Cydalima perspectalis)* in North America using genome-wide SNP data complemented by mitochondrial DNA and morphological analyses. Our analyses revealed substantial variation in genetic diversity, population structure, genetic differentiation, allele sharing, and demographic history among invasive populations in Europe and North America. Notably, the combined genomic and mitochondrial evidence identified at least two distinct invasion signatures in North America, with Delaware consistently differentiated from the remaining sampled North American populations. These patterns indicate that the contemporary North American invasion is unlikely to reflect a single, genetically homogeneous introduction and instead may involve multiple introduction sources and/or subsequent population admixing. Together, these findings provide a genomic framework for understanding the establishment and spread of *C. perspectalis* in North America and offer information that can be used to improve invasion surveillance, identify potential introduction pathways, and inform targeted containment and long-term management strategies.

### A genetically complex invasion history

The genetic structure observed across *C. perspectalis* populations provides several lines of evidence for a complex invasion history in North America. Broad-scale patterns from PCA and ADMIXTURE, together with the finer-scale relationships identified by fineRADstructure, revealed that North American populations are not genetically homogeneous and do not form a simple geographic continuum of differentiation. Instead, populations differed substantially in their genomic affinities, with some geographically distant populations showing closer genetic relationships than expected from geographic distance alone. Pairwise F_ST_ analyses similarly revealed pronounced heterogeneity in genetic differentiation, while Dsuite ABBA-BABA analysis identified significant asymmetric allele sharing among several North American population combinations. The agreement among these independent approaches suggests that the observed genetic structure reflects key differences in population history rather than variation attributable to a single analytical method.

The mitochondrial data provide an additional perspective on this pattern. Most sampled North American populations, including Canada, shared the same HTB *COI* haplogroup lineage, whereas Delaware shared the HTA lineage with the Croatian laboratory colony. Thus, mitochondrial variation suggests a predominant maternal lineage across much of the sampled North American invasion, while the nuclear genome reveals substantially greater differentiation and complex patterns of allele sharing. The contrast between mitochondrial and nuclear patterns is informative because it indicates that the contemporary genetic composition of North American populations cannot be fully described by a single mitochondrial lineage or a simple geographic expansion model. Rather, the data are consistent with a history involving differentiated introductions, subsequent admixture or secondary contact, and/or demographic processes acting after establishment.

### Delaware represents a distinct invasion signature

Among the North American populations examined, Delaware showed the most distinctive combination of mitochondrial, nuclear genomic, and demographic characteristics. Unlike most sampled North American populations, which carried the HTB *COI* lineage, Delaware shared the HTA mitochondrial haplotype group with the Croatian laboratory colony. This shared mitochondrial lineage provides an independent indication of a maternal connection between Delaware and the European reference population. Although mitochondrial similarity alone cannot establish a direct source–recipient relationship, its occurrence in Delaware but not in the other sampled North American populations is notable and is consistent with a distinct introduction history for the Delaware population.

The nuclear genomic data further distinguish Delaware from the broader North American invasion. Delaware maintained relatively high nucleotide diversity and relatively high observed heterozygosity, while exhibiting low inbreeding. This combination contrasts with several other North American populations, including Massachusetts, Michigan, Ohio, Virginia, and West Virginia, which showed lower heterozygosity and substantially higher levels of inbreeding. High genetic diversity in an introduced population can result from a relatively large founding population, repeated introductions from genetically differentiated sources, admixture among previously differentiated lineages, or continued gene flow following establishment (Dlugosch and Parker 2008). Thus, the elevated diversity observed in Delaware may reflect a more complex introduction or establishment history than that experienced by populations that underwent stronger founder effects and genetic drift.

ABBA-BABA analyses provide further evidence that Delaware occupies a distinctive position within the North American invasion. Delaware repeatedly occurred in significant P2–P3 combinations, indicating excess allele sharing between Delaware and other North American populations relative to the corresponding P1 population. These relationships are consistent with shared ancestry, admixture, or historical gene flow and suggest that the Delaware population may have contributed genetically to, or subsequently interacted with, other introduced populations. Importantly, however, the direction of these relationships cannot be inferred from the D-statistic alone. Therefore, the combined evidence supports Delaware as a genetically distinctive population and a potentially important component of the North American invasion history, but does not by itself establish Delaware as the original point of introduction or a definitive source population for subsequent spread.

From an invasion-management perspective, the distinctive genetic signature of Delaware is important because it demonstrates that geographically established populations may represent different introduction histories even within a relatively young invasion. Delaware therefore represents a population that warrants particular attention in future sampling and surveillance efforts. Expanded sampling of populations surrounding Delaware, together with specimens from nurseries, commercial plant-production facilities, and additional European source regions, would help determine whether the Delaware lineage represents an independent introduction, an early-established population that subsequently expanded, or a population formed through repeated introductions and admixture.

### Regional differentiation suggests multiple North American invasion histories

Beyond the distinctive signal observed in Delaware, the genome-wide analyses revealed additional geographic patterns among North American populations. Canada, New York, Massachusetts, Michigan, and Ohio showed relatively close genetic relationships across multiple population structure analyses, suggesting a shared or closely related invasion history. This pattern is particularly notable because independent historical and trade-based evidence indicates that the establishment of *C. perspectalis* in U.S. populations near the Canadian border was likely facilitated by movement of infested boxwood plants from Canada (APHIS 2021). The concordance between this documented pathway and the genomic similarity observed among Canadian and several U.S. populations provides support for a Canadian-associated invasion pathway contributing to the establishment and spread of *C. perspectalis* in North America.

The genetic similarity among Canada, New York, Massachusetts, Michigan, and Ohio is consistent with expansion or repeated human-mediated movement of a related invasion lineage following its establishment in North America. Michigan and Ohio showed particularly similar genomic profiles and demographic trajectories. These parallel demographic patterns are consistent with a shared population history and may reflect expansion of a common introduced lineage followed by subsequent regional contraction or fragmentation. New York and Massachusetts also showed relatively low contemporary genetic diversity, which may reflect founder effects and subsequent genetic drift following establishment. Although the precise demographic timing inferred by GONE should be interpreted cautiously considering their <10 generations in North America, the concordance among population structure, genetic diversity, and demographic trajectories supports a common or closely related invasion history among these populations.

The Canadian-associated pattern is also consistent with the mitochondrial results. Canada and most of the sampled U.S. populations shared the HTB COI lineage, indicating that the predominant mitochondrial lineage in the North American invasion extends across both sides of the U.S.–Canadian border. Independent field observations provide additional evidence that movement across the Canada - U.S. border may contribute to the spread of BTM. During a 2024 BTM trapping survey in New York, positive traps were detected along the Lake Ontario and Lake Erie shorelines, including locations without nearby boxwood, providing independent evidence of BTM dispersal across or along the Great Lakes **(Supplementary Fig. 1)**. Such observations are consistent with the capacity of adult moths to disperse beyond immediate host-plant patches and suggest that geographic barriers such as the Great Lakes may not prevent movement between Canadian and U.S. populations. However, this interpretation does not imply that all North American populations originated from Canada. Rather, it suggests that Canadian populations may have served as an important source for at least part of the subsequent establishment and spread of *C. perspectalis* in the United States.

In contrast, Virginia and West Virginia showed a distinct genomic signal among the non-Delaware North American populations. Despite their geographic proximity, these populations did not show the same close association with the Canada–New York–Massachusetts–Michigan– Ohio group. This differentiation suggests that these Appalachian populations may represent an additional introduction history or a population lineage that became differentiated following establishment. Their distinct genomic composition could reflect an independent introduction, limited connectivity with the Canadian-associated lineage, or subsequent genetic drift within geographically isolated populations. Distinguishing among these possibilities will require additional sampling from populations throughout the Appalachian region and from potential source populations.

### Trade and transportation as likely invasion pathways

The broader role of transportation networks in facilitating biological invasions is well documented. Historical analyses of U.S. port-interception records have found that 73% of pest interceptions between 1984 and 2000 occurred at international airports, with 62% of intercepted pests associated with baggage, most of which were insects (McCullough et al. 2006). Contemporary border-security activities demonstrate that these pathways remain highly active. In fiscal year 2023, USDA Plant Protection and Quarantine (PPQ) processed and identified more than 92,000 pest interceptions, approximately 45,000 of which were considered quarantine-significant, while U.S. Customs and Border Protection (CBP) agriculture specialists continue to intercept hundreds of agricultural pests through U.S. ports of entry (USDA-APHIS 2023). These patterns emphasize the continuing importance of international trade and transportation as pathways for the introduction and redistribution of non-native organisms.

Boxwood plants have a long history of cultivation and trade in North America, providing an established network through which box tree moth could be transported and introduced into new regions (Coyle et al. 2022; Wiesner 2023). Today, the United States remains a major producer of boxwood, with substantial commercial production concentrated in several states, including Ohio, Oregon, and California (Hall et al. 2021). At the same time, the United States has historically imported boxwood from Canada, creating an additional pathway for the movement of both plants and associated pests (Gao et al. 2023; Seehausen et al. 2024b).

The genetic evidence for multiple invasion signatures identified in this study is therefore consistent with the well-established movement of boxwoods through domestic and international trade networks. Such pathways can facilitate both long-distance dispersal and repeated introduction into geographically separated regions. In particular, the close genetic relationship observed among Canada, New York, Massachusetts, Michigan, and Ohio is consistent with a Canadian-associated invasion pathway, given independent evidence that movement of potentially infested boxwood from Canada contributed to the establishment of box tree moth in U.S. states sharing the Canadian border. Conversely, the distinct genetic signature of Delaware and its shared mitochondrial lineage with the Croatian laboratory colony suggest that at least some North American populations may have a different introduction history. These contrasting patterns highlight how plant trade can generate a geographically complex invasion landscape in which different populations may arise through independent introduction events.

Repeated introductions through ornamental plant shipments can also increase propagule pressure and introduce genetically differentiated individuals, potentially increasing genetic diversity and facilitating establishment (Cassey et al. 2018; Bertelsmeier et al. 2018b). The relatively high genetic diversity observed in Delaware, combined with its distinct mitochondrial and nuclear genomic signatures, is consistent with such a scenario, although the available data cannot determine whether Delaware resulted from a single introduction of a genetically diverse source population or from multiple introductions. More broadly, the correspondence between known plant-trade pathways and the genetic structure observed in *C. perspectalis* highlights the value of integrating genomic surveillance with trade and transportation information. Such integration could help identify high-risk pathways, prioritize surveillance locations, and distinguish newly introduced populations from secondary spread of established North American lineages.

### Implications of this study for management of box tree moth in North America

Despite its rapid spread and the threat posed by *C. perspectalis*, boxwood remains an important component of the North American nursery and ornamental horticulture industry because of its substantial economic value and distinctive horticultural characteristics. Boxwood tolerates a broad range of environmental conditions, withstands frequent pruning, provides year-round evergreen structure, and is relatively resistant to deer browsing (Boggs et al. 2025; Gardenia 2026). Because alternative plant species may not readily reproduce this combination of characteristics, large-scale replacement of boxwood is unlikely to be economically or practically feasible. Management of box tree moth must therefore balance protection of the nursery industry and ornamental landscapes with efforts to limit further establishment and spread of this invasive pest.

A previous study using climatic suitability models indicates that substantial portions of North America remain environmentally suitable for continued expansion of *C. perspectalis* (Seehausen et al. 2024b). Combined with its current distribution and the evidence for multiple introduction signatures identified in this study, these patterns suggest that management strategies focused solely on eradication may become increasingly difficult as the invasion progresses. Instead, long-term management should emphasize early detection, containment of newly established populations, mitigation of economic impacts, development of resistant or less susceptible boxwood cultivars, and implementation of effective biological control strategies.

The distinction between European and North American host landscapes may also influence its future invasion dynamics. In Europe, native wild boxwoods can form relatively continuous forest habitats that facilitate landscape-level dispersal of box tree moth between urban and natural areas (Seehausen et al. 2024). North America lacks native boxwood species, and host plants are primarily concentrated in managed landscapes, nurseries, gardens, and other human-associated environments. Consequently, boxwood in North America may occur as spatially discrete host patches or “islands” rather than as continuous host corridors. This fragmented distribution could create opportunities for targeted surveillance and containment, particularly around nurseries and other locations where large numbers of susceptible plants are concentrated. Strategic monitoring of nurseries, commercial plant-production facilities, transportation corridors, and other points associated with plant movement may therefore provide an effective means of detecting new introductions before populations become widely established.

Our results further demonstrate why such surveillance should consider the possibility of multiple introduction sources. The distinct genetic signatures observed among Delaware, the Canada– New York–Massachusetts–Michigan–Ohio group, and the Virginia–West Virginia populations indicate that newly detected populations may not necessarily represent simple local expansion from the nearest established population. Instead, such genomic characterizations can help determine whether newly detected populations are genetically consistent with previously characterized lineages or represent potentially independent introductions. This distinction is important for management because independent introductions may require different pathway investigations and biosecurity responses than secondary spread from an established population.

Our findings therefore support greater integration of genomic surveillance, international biosecurity, plant-trade monitoring, and coordinated cross-border management. Existing domestic quarantine measures implemented by the U.S. Department of Agriculture provide an important component of containment, but genomic tools can complement these efforts by identifying potential source populations, reconstructing introduction pathways, and distinguishing newly introduced populations from secondary spread. Establishing a reference genomic database of North American and potential source populations would allow newly detected specimens to be compared against known invasion signatures and could provide actionable information for pathway tracing and risk assessment.

Understanding invasion pathways may also facilitate future studies of host susceptibility by linking genetically distinct BTM populations to variation in their interactions with boxwood. Identifying whether particular invasion lineages differ in host use or susceptibility could help isolate and evaluate resistant boxwood lineages and, ultimately, inform containment and long-term management strategies.

## Author contributions

SL, SE, and HY conceived the idea; AB, JA, CL, LS, and YW performed the fieldwork and collected samples; AB, TC, and SL carried out the laboratory experiments and data analysis. AB, YW, and SL wrote the manuscript with input from all authors. All authors read and approved the manuscript.

## Data availability

All short-read genome sequencing data and mtDNA sequences have been submitted to the National Center for Biotechnology Information (NCBI) under BioProject Accession number PRJNA1524226.

## Acknowledgement

We thank the members of the Lamichhaney Laboratory (Carter Henry and Dominique Costarella) at Kent State University for their assistance with specimen collection, laboratory work, morphological measurements, and data processing. We thank collaborators and colleagues from the Ohio Department of Agriculture (ODA), Michigan Department of Agriculture and Rural Development (M-DARD) Plant Health Section, New York State Department of Agriculture and Markets, and Forest Pest Methods Laboratory (FPML, USDA) who assisted with collection and provision of box tree moth specimens from North America. We also thank Prof. Aibin Zhan for assisting us in collecting specimens of the native population of box tree moth from East Asia. We also thank the Kent State University research facilities and computational resources used for genomic data processing and analysis. We also thank the U.S. Department of Agriculture (USDA) and other regulatory and scientific agencies whose publicly available information on box tree moth detections, plant trade, and pest interceptions contributed to our understanding of invasion pathways and management context.

## Ethics approval

Not applicable.

## Consent to participate

Not applicable.

## Consent for publication

Not applicable.

