## Supplementary Materials for "Multiple introductions shaped genomic diversity and demographic history of invasive box tree moth (*Cydalima perspectalis*) populations in North America"

**Supplementary Table 1.** D-statistic (ABBA–BABA) tests of allele sharing among *Cydalima perspectalis* populations. The table summarizes D-statistic tests performed using the indicated outgroup and population assignments. Outgroup indicates the population used as the phylogenetic outgroup; P1 and P2 represent the two focal populations being compared for differential allele sharing, and P3 represents the putative source or recipient population. The D statistic measures an excess of ABBA versus BABA allele patterns, with positive and negative values indicating asymmetric allele sharing between P2 and P3 relative to P1. The Z-score was calculated using a block-jackknife procedure to assess the significance of the D statistic. p-value represents the corresponding statistical significance. The f4-ratio provides an estimate of the proportion of ancestry in the target population attributable to gene flow from the putative donor population. Tests were interpreted as evidence of significant asymmetric allele sharing when $|Z| \geq 3$.

| **Outgroup** | **P1** | **P2** | **P3** | **D statistic** | **Z-score** | **p-value** | **f4-ratio** |
|---|---|---|---|---|---|---|---|
| Croatia | Michigan | Ohio | Delaware | 0.69712 | 13.3301 | <1e-10 | 0.37257 |
| Croatia | Michigan | Newyork | Delaware | 0.679953 | 10.9743 | <1e-10 | 0.361849 |
| Croatia | Michigan | Newyork | Switzerland | 0.776374 | 10.1985 | <1e-10 | 0.53534 |
| Croatia | Michigan | Ohio | Switzerland | 0.685289 | 8.63417 | <1e-10 | 0.480015 |
| Croatia | Michigan | Virginia | Delaware | 0.499371 | 8.52719 | <1e-10 | 0.145543 |
| Croatia | Michigan | WestVirginia | Switzerland | 0.488123 | 8.27382 | <1e-10 | 0.291955 |
| Croatia | Michigan | Virginia | Switzerland | 0.56306 | 7.41677 | <1e-10 | 0.273909 |
| Croatia | Massachusetts | Newyork | Switzerland | 0.674675 | 7.38656 | <1e-10 | 0.44065 |
| Croatia | Michigan | WestVirginia | Delaware | 0.489267 | 6.88775 | <1e-10 | 0.187183 |
| Croatia | Canada | Newyork | Switzerland | 0.486537 | 5.77985 | 3.74E-09 | 0.325489 |
| Croatia | Michigan | Massachusetts | Delaware | 0.255543 | 5.42407 | 2.91E-08 | 0.0995308 |
| Croatia | Massachusetts | Ohio | Canada | 0.372251 | 4.71469 | 1.21E-06 | 2.02267 |
| Croatia | Canada | Newyork | Delaware | 0.348051 | 4.7069 | 1.26E-06 | 0.174848 |
| Croatia | Massachusetts | Newyork | Delaware | 0.501427 | 4.60526 | 2.06E-06 | 0.205812 |
| Croatia | Canada | WestVirginia | Switzerland | 0.340366 | 4.50752 | 3.28E-06 | 0.240282 |
| Croatia | Michigan | Massachusetts | Switzerland | 0.296254 | 4.31627 | 7.93E-06 | 0.146461 |
| Croatia | Canada | WestVirginia | Delaware | 0.324471 | 4.04241 | 2.65E-05 | 0.138379 |
| Croatia | Canada | Virginia | Switzerland | 0.328545 | 4.03892 | 2.68E-05 | 0.192591 |
| Croatia | Canada | Virginia | Delaware | 0.28299 | 4.03171 | 2.77E-05 | 0.0908613 |
| Croatia | Massachusetts | Newyork | Canada | 0.330629 | 3.78624 | 7.65E-05 | 0 |
| Croatia | Massachusetts | Ohio | Delaware | 0.422592 | 3.72747 | 9.67E-05 | 0.186773 |
| Croatia | Canada | Ohio | Delaware | 0.29658 | 3.57342 | 0.00017617 | 0.151479 |
| Croatia | Canada | Massachusetts | Delaware | 0.159775 | 3.45932 | 0.00027077 | 0.072722 |
| Croatia | Michigan | Massachusetts | Canada | 0.109475 | 3.3527 | 0.00040013 | 2.87032 |

| | | | | | | | |
|---|---|---|---|---|---|---|---|
| Croatia | Canada | Ohio | Switzerland | 0.25467 | 3.00999 | 0.0013063 | 0.193971 |
| Switzerland | Newyork | Michigan | Croatia | 0.776374 | 10.1985 | <1e-10 | 0.237599 |
| Switzerland | Ohio | Michigan | Croatia | 0.685289 | 8.63417 | <1e-10 | 0.224593 |
| Switzerland | WestVirginia | Michigan | Croatia | 0.488123 | 8.27382 | <1e-10 | 0.106763 |
| Switzerland | Virginia | Michigan | Croatia | 0.56306 | 7.41677 | <1e-10 | 0.09783 |
| Switzerland | Newyork | Massachusetts | Croatia | 0.674675 | 7.38656 | <1e-10 | 0.13588 |
| Switzerland | Newyork | Canada | Croatia | 0.486537 | 5.77985 | 3.74E-09 | 0.106582 |
| Switzerland | WestVirginia | Canada | Croatia | 0.340366 | 4.50752 | 3.28E-06 | 0.0694813 |
| Switzerland | Massachusetts | Michigan | Croatia | 0.296254 | 4.31627 | 7.93E-06 | 0.057832 |
| Switzerland | Virginia | Canada | Croatia | 0.328545 | 4.03892 | 2.68E-05 | 0.0585798 |
| Switzerland | Michigan | Massachusetts | Newyork | 0.307882 | 3.15657 | 0.00079817 | 0.545659 |
| Switzerland | Michigan | Massachusetts | Ohio | 0.306408 | 3.09244 | 0.00099261 | 0.493357 |
| Switzerland | Ohio | Canada | Croatia | 0.25467 | 3.00999 | 0.0013063 | 0.0646076 |

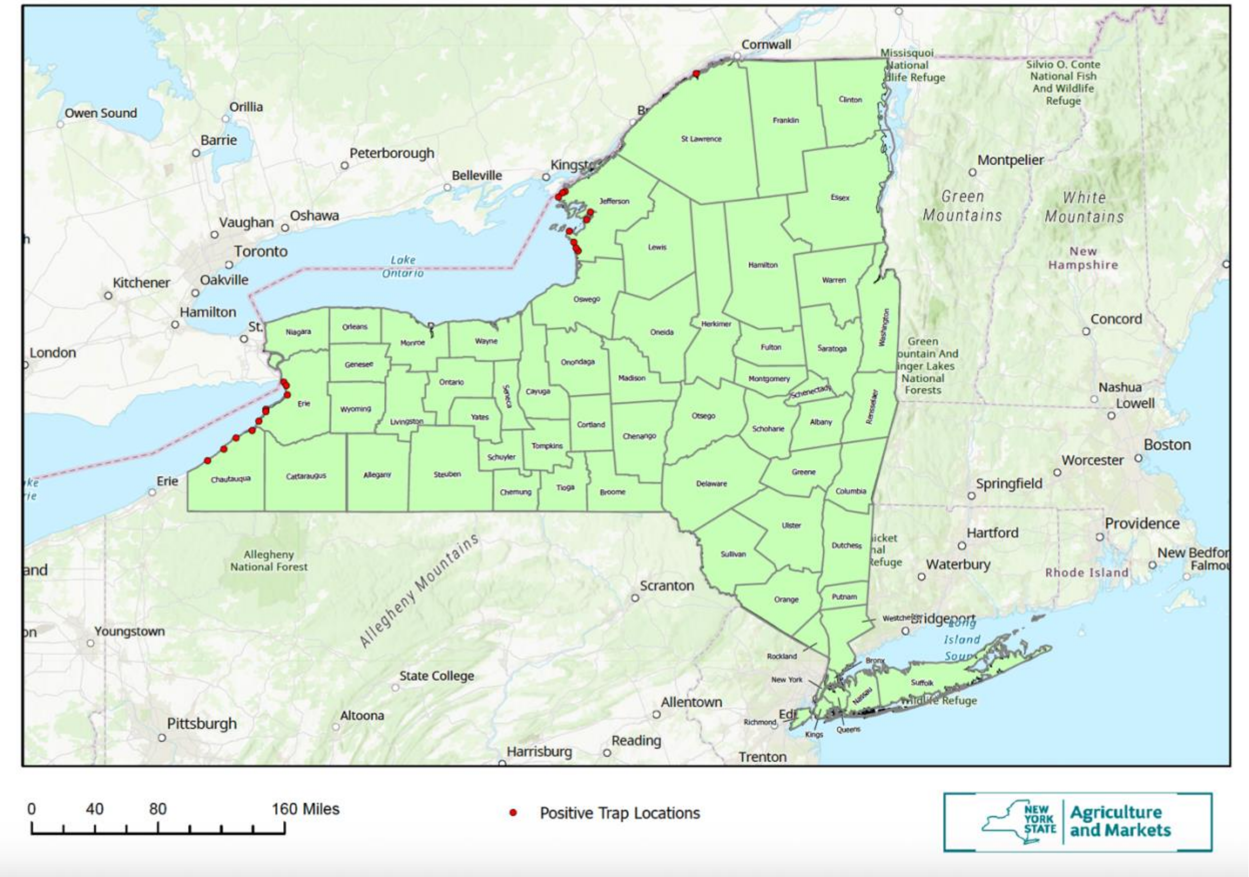


**Supplementary Figure 1:** Box tree moth *(Cydalima perspectalis)* detections along the Lake Ontario and Lake Erie shorelines in New York during the 2024 trapping survey © New York State Department of Agriculture and Markets. The map shows the locations of traps that tested positive for *C. perspectalis* during the 2024 survey. Positive detections occurred along the Lake Ontario and Lake Erie shorelines, including locations where no boxwood host plants were present in the immediate vicinity.
